# Design of a Companion Bioinformatic Tool to detect the emergence and geographical distribution of SARS-CoV-2 Spike protein genetic variants

**DOI:** 10.1101/2020.06.22.133355

**Authors:** Alice Massacci, Eleonora Sperandio, Lorenzo D’Ambrosio, Mariano Maffei, Fabio Palombo, Luigi Aurisicchio, Gennaro Ciliberto, Matteo Pallocca

## Abstract

**Background:** Tracking the genetic variability of Severe Acute Respiratory Syndrome CoronaVirus 2 (SARS-CoV-2) is a crucial challenge. Mainly to identify target sequences in order to generate robust vaccines and neutralizing monoclonal antibodies, but also to track viral genetic temporal and geographic evolution and to mine for variants associated with reduced or increased disease severity. Several online tools and bioinformatic phylogenetic analyses have been released, but the main interest lies in the Spike protein, which is the pivotal element of current vaccine design, and in the Receptor Binding Domain, that accounts for most of the neutralizing the antibody activity.

**Methods:** Here, we present an open-source bioinformatic protocol, and a web portal focused on SARS-CoV-2 single mutations and minimal consensus sequence building as a companion vaccine design tool. Furthermore, we provide immunogenomic analyses to understand the impact of the most frequent RBD variations.

**Results:** Results on the whole GISAID sequence dataset at the time of the writing (October 2020) reveals an emerging mutation, S477N, located on the central part of the Spike protein Receptor Binding Domain, the Receptor Binding Motif. Immunogenomic analyses revealed some variation in mutated epitope MHC compatibility, T-cell recognition, and B-cell epitope probability for most frequent human HLAs.

**Conclusions:** This work provides a framework able to track down SARS-CoV-2 genomic variability.

## Background

With more than 50 million infected people and 1.3 million deaths, SARS-CoV-2 is likely to have found a reservoir in the human population, as demonstrated by the current 2020 boreal autumn outbreak. To prevent the viral spread, an effective vaccine is needed.

More than 100 different vaccines are in development worldwide, including China, India, the USA, and Europe. Most, if not all of them, target the Spike protein, the viral product able to bind the human receptor angiotensin-converting enzyme 2 (ACE2). These designs use different formulations or platforms, such as a vectored vaccine or nucleic acids, RNA, and DNA. The Spike protein is the best candidate for historical reasons[1] and recent preclinical evidence in non-human primates [2].

One of the main challenges is that from preclinical vaccine testing to the first-in-man trial to confirmatory trials, regulatory approval, and large-scale vaccine distribution, from 6 to 12 months will elapse. During this time frame, will the Spike protein of the circulating virus mutate? If so, are these genetic variants going to escape protective immune responses induced by the vaccine? In this paper, we try to provide a bioinformatic framework to address this potential issue.

A few preliminary reports already described both the main differences with the SARS-CoV virus [3] and the variant landscape of SARS-CoV-2 in different clades [4–6] Both pointed out the limited presence of functional mutations in critical regions of the genome; some groups also released open bioinformatic web applications to browse the virus variants [7]. Nevertheless, some other reports stressed a possible selection pressure on the D614G variant, the only frequent variation of the spike protein [8], recently providing in-vivo evidence of its increased fitness [9] ⍰. A bioinformatic analysis of SARS-CoV-2 epitopes showed high homology with SARS-CoV [10]⍰and, thus, characterized many possible B- and T-cell epitopes, providing an *in vivo* characterization of the virus proteins that are most targeted by T cells, confirming the Spike protein to be the first region of interest [11]⍰. Of note, it was recently reported from a study involving the plasma of 650 SARS-CoV-2 exposed patients that 90% of the serum or plasma activity targets the Receptor Binding Domain (RBD) of the Spike protein, the central point of SARS-CoV-2/ACE2 contact.

To provide an evidence-based approach, we present a freely available protocol to enable all the bioinformatic community to dissect SARS-CoV-2 genomic variability and its functional impact on DNA vaccines and an openly accessible web portal to browse the current most frequent viral mutations. This framework can support researchers to automatically mine data useful for vaccine design from a pool of viral sequences and a target protein or domain of interest: a consensus sequence, in terms of average or minimal identity and a list of mutations with their absolute frequencies.

## Methods

### Bioinformatic framework

Following the overall workflow (Figure 1A), a multi-FASTA file of viral sequences is aligned against the Wuhan strain (NC_045512.2) using NUCmer from the MUMmer package [12]. Filters used to fetch the sequences analyzed in this manuscript were Host=Human, Virus=hCov-19, flagged “complete”, “high coverage,” and “low coverage exclusion”. Nucmer generates a delta encoded alignment file, which is then parsed using the show-snps utility. This produces a catalog of all the SNPs and indels internal to the alignments contained in the delta encoded file. Show-snps output is then converted into standard VCF format using code adapted from [13].

**Figure 1:**
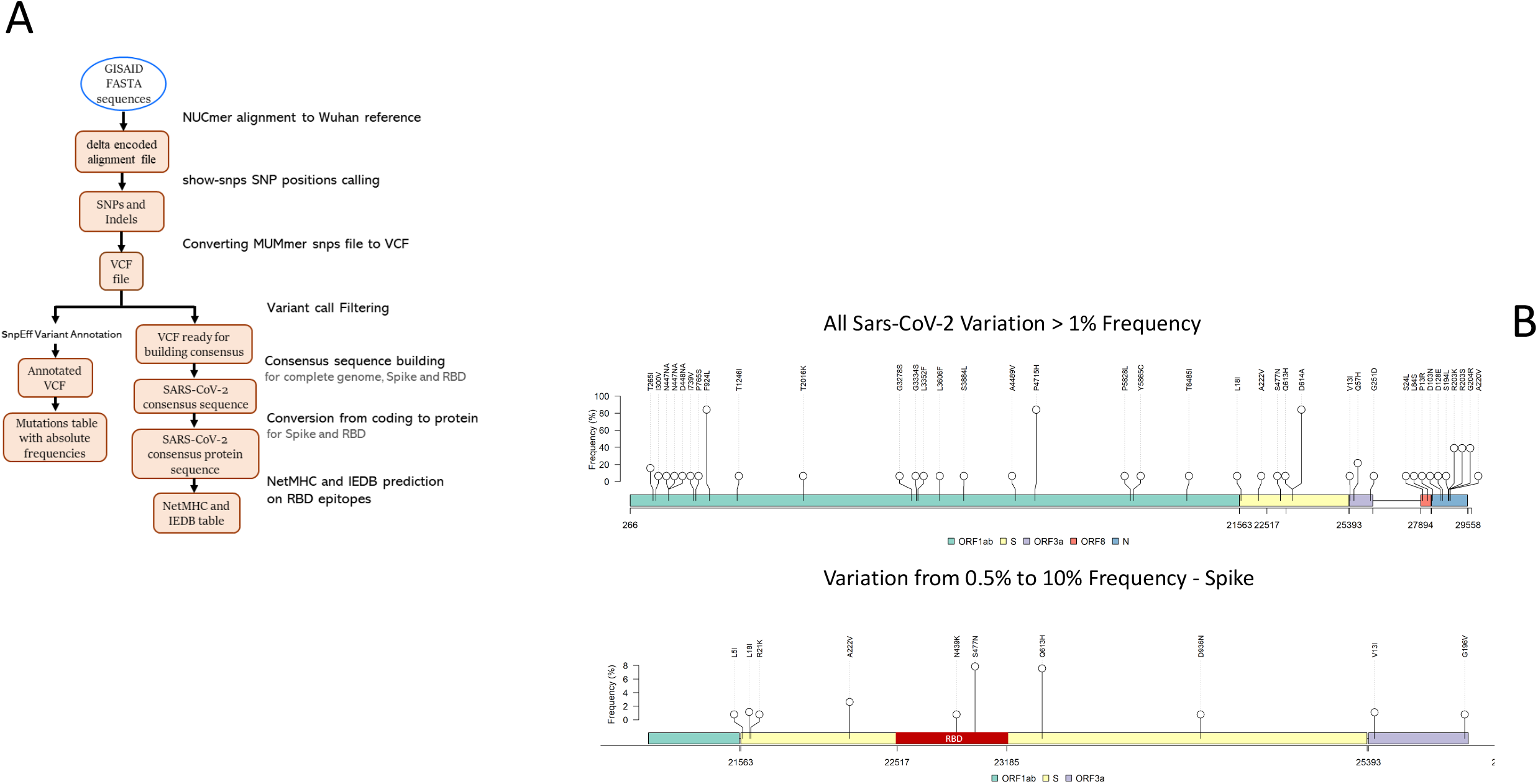
A: Workflow of the Covid-Miner package. The execution is fully automatized and encapsulated in a Docker container. B-C: Lollipop representation of SARS-CoV-2 variants at 1% frequency for the whole SARS-CoV-2 genome and from 0.1% to 10% for the Spike protein locus, showing the most frequent RBD variants at codon 447 (7%) and 439 (0.7%). D: Barplot from occurrences of most frequent RBD variations.

At this point, it is necessary to annotate said variants in terms of functional effect and penetration in the viral population while producing a consensus sequence that represents the most frequent allele for each position. The *consensus* command from bcftools builds the said consensus from the filtered VCF file. In addition to the consensus for the complete viral genome, we build the consensus sequence separately for the spike protein and RBD. Moreover, a consensus of minimal identity is constructed, applying the N character at variant sites instead of the most frequent base. Such consensus of minimal identity, by showing which residues are conserved and which residues are variable, would help in the design of a vaccine.

Variant annotation is employed via the *snpEff* package [14] that embeds the NC_045512.2 genome assembly annotation in its standard package. A human-readable (and easily parsable) table is formatted thanks to the *SnpSift* jar package [15] of *snpEff*. 2D visualizations of the variants on the viral genome are generated via the LolliPlot function of the trackVignette′s R package [16].

The whole toolset and test sequence files have been included in a Docker image, enabling users to run the whole pipeline with just one command (e.g., *docker run covid-miner sequences*.*fasta*). The whole workflow, with sample test data and Dockerfile for portability and reproducibility purposes, is available on https://gitlab.com/bioinfo-ire-release/covid-miner.

Downstream immunogenomic analyses were carried out by NetMHC for Class 1 recognition [17], IEDB for T-cell immunogenicity [18] and BepiPred for B-cell epitopes [19]. The web portal has been implemented in Angular (version 10.1) over the Bootstrap CSS framework (version 4.5) by leveraging on D3js library (version 6.2) for the graphical representations. The backend is written in Python language by adopting the Flask web framework (version 1.1.2).

## Results

### Analysis results on 93930 SARS-CoV-2 sequences

We extracted 93930 high-quality sequences (available on October 2, 2020) from the the EpiCov™ section of the GISAID portal [20] that acts as a worldwide repository of viral isolates. Every viral isolate contains zero or more variants with respect to the Wuhan strain. The fully annotated table of 20640 variants is available in Table S1.

Considering the total number of sequences, the most frequent Spike protein variation is confirmed to be D614G (Figure 1B). Next, there are a few variants with a frequency of 2-7% that are slowly rising in the viral populations, namely S477N, Q613H and A222V. The S477N mutation lies in the RBD locus, while all the other RBD variants are below the 1% penetrance threshold (Figure 1C, Table S2).

We asked whether the RBD variants are strongly associated with a certain geographical region, to this purpose we propose to measure “mutations per thousand isolates”, that is

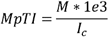

with *M* as the absolute mutation count and *I*_*c*_ the total isolates for that country. This normalization smooths out the inter-country variability, but the resolution bias remains due to the high variability in isolate sequencing among countries, changing in order of magnitude from thousands to dozens. This bias will cause rare mutations to be hidden in countries with a few associated isolates, while the associations with highly frequent outliers will remain more robust. The geographical distribution shows how the S477N variant is strongly rooted in Australia, and the N439K is associated with clusters starting from the United Kingdom (Scotland). However, none of the most frequent variants are uniquely associated with one country as expected from the worldwide virus distribution, and clusters of co-occurring variants lie mostly in countries with the highest number of available sequences (i.e., USA, England) (Table S2, Figure 2A-B). In order to better understand the evolution of variants over time and space, we tracked the location of all the source isolates carrying these two variations (Figure 2C). The S477N has been firstly identified in Colombia and is harbored in more than 60% of the isolates sequenced in Australia from June 2020. On the other hand, the N439K is dominating the isolates landscape in Ireland and England from August 2020.

**Figure 2:**
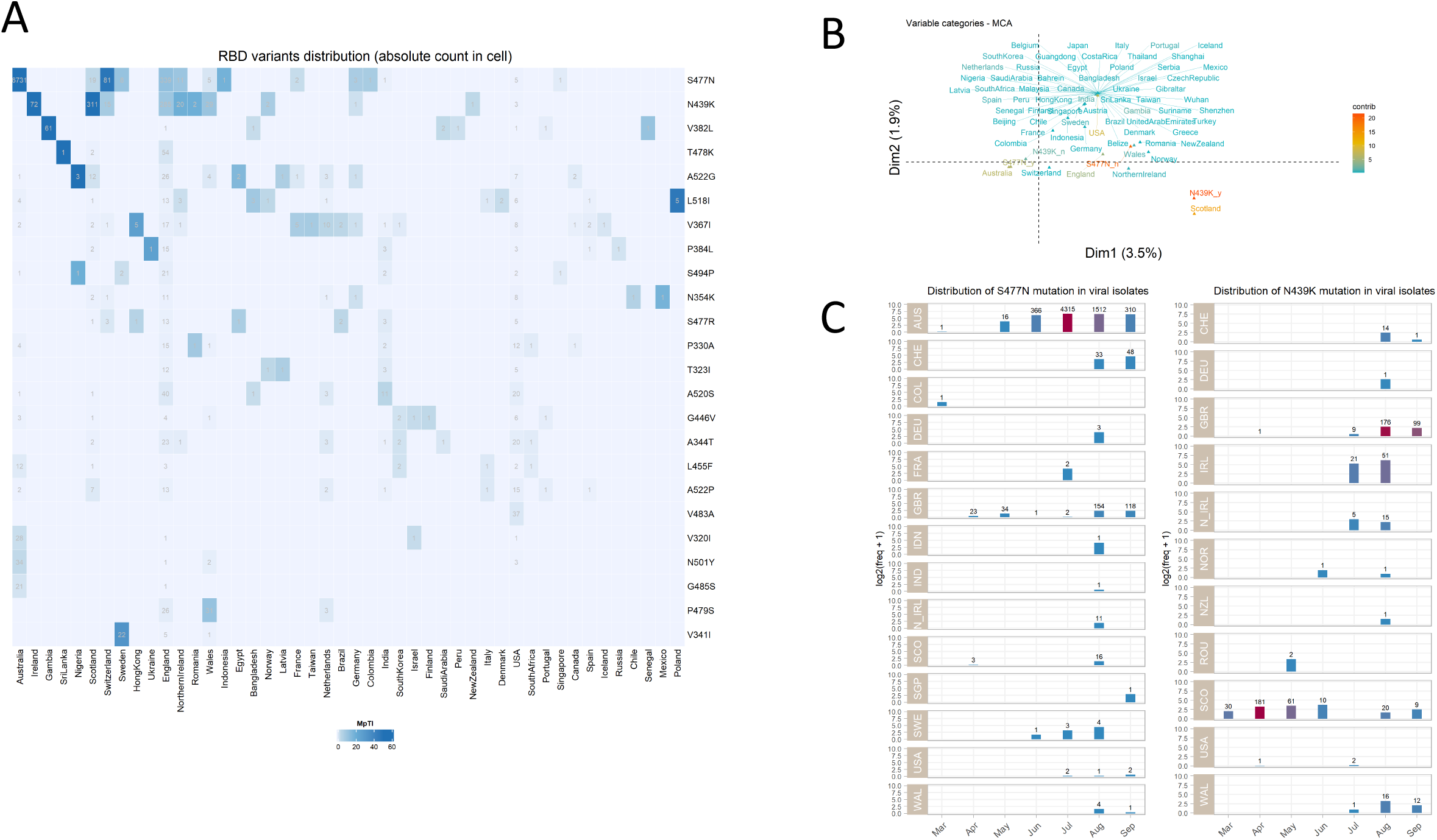
A: Heatmap of Receptor Binding Domain variants geographical distribution by normalized frequency. Countries with low occurrences of RBD variants (less than 5) were removed. B: Multiple Correspondence Analysis (MCA) of Country-RBD variant association. C: Evolution of S447N and N439K in space and time. Y-axis represents the relative frequency of that mutation in every country isolate set for each month. X-axis: months in 2020, as annotated in GISAID sequences. Every bar is annotated with absolute counts.

### Immunogenomic Analysis

These novel RBD variants may have several biological and putatively clinical impacts on the virus functions. For instance, every protein-coding variant changes several epitope sequences presented on the human cell′s surface from the Major Histocompatibility Complex (MHC). This, in turn, can have an impact on the human immune system recognition by T and B-cells. To shed light on these processes, we computationally modeled the immunological impact of SARS-CoV-2 epitopes in terms of (1) MHC Class 1 presentation of antigens (2) T-cell immunogenicity (3) B-cell epitope prediction.

Considering the only RBD mutations above 0.1% frequency, we performed MHC class 1 binding prediction for both the wild-type (Wuhan strain) and the mutated strain, choosing the most frequent HLAs in public databases [21]. The software generated predictions for all possible 9-mers resulting from the full RBD sequence, and we computed how *binders* were calculated for all the considered HLAs. Out of 18 predictions, only one epitope had binding affinity in mutated or wild-type, with no change in binding affinity caused by the mutation(Table S3).

| Mutation | WT | Mutant | Wild-Type binders | Mutated binders |
| --- | --- | --- | --- | --- |
| S477N | IYQAGSTPC | IYQAGNTPC | 1 | 1 |

Then, we asked whether these mutations had a significant impact on T- and B-cell recognition, and we ranked them via the Immune Epitope Database and analysis resource (IEDB) [18]. When considering T-cells with the class 1 immunogenicity tool [22]⍰:, 54/189 (29%) epitopes showed a negative, non-immunogenic score in both wild-type and mutated forms. When focusing on the predicted class 1 binders for the same epitope/HLA combination in mutated peptides, IYQAGNTPC (S477N) shows increased immunogenicity for all considered HLAs (Figure 3A). These results point out to a putative variability in T-cell response mediated by these mutations.

**Figure 3:**
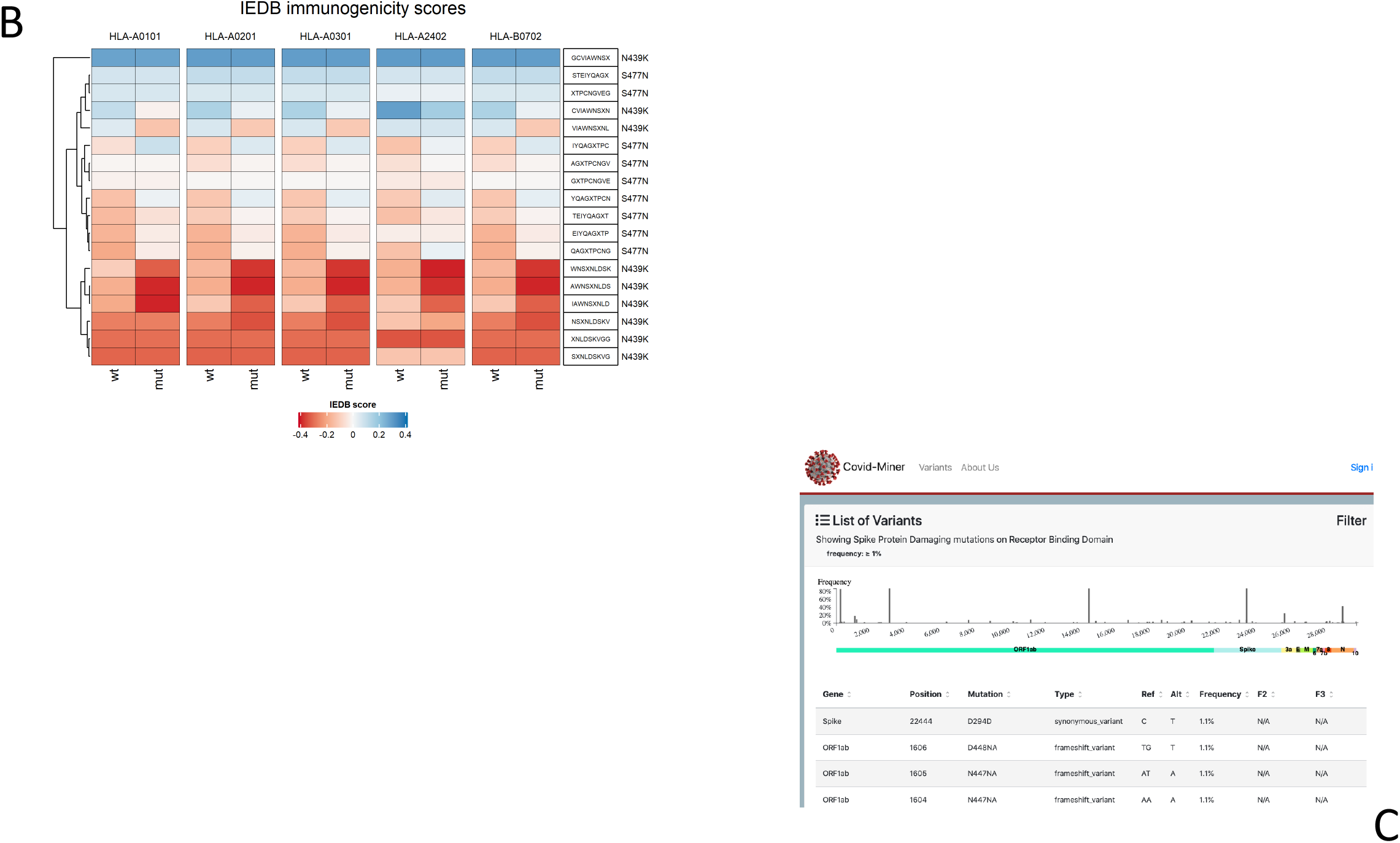
A: Heatmap representing T-cell immunogenicity scores for wild-type and mutated RBD epitopes. B: Screenshot of covid-miner web portal homepage. C: Donut plot of RBD mutational impact of overall sequences and September 2020 sequences. D: Location of the most frequent RBD mutated amino acids N439 and S477.

Testing epitopes for putative B-cell recognition remains an analytical challenge; only a few algorithms have been developed for this purpose [23]. When focusing to all the mutated epitopes caused by S477N and N439K, none cause a shift of the amino acid exposition, as they are both predicted to be in the *exposed* status. The overall Epitope score, that takes into account a variable amino acid window surrounding the mutation, is slightly increased, from 0.535/0.516 in the WT sequence to 0.561 and 0.548 for S477N and N439K, respectively (Table S5).

### Covid-Miner Data Portal

In order to share the results with a wider public, we created a frontend portal for the analysis results, freely accessible at https://covid-miner.ifo.gov.it. The web app features two sections; a *Variant* section is dedicated to browse and visualize the most frequent viral mutations over the genome, and a *Geographical* distribution heatmap that displays the association among countries and the most frequent RBD mutations. The home page displays the amount of wild type / mutated RBD variants as a main summarizing figure (Figure 3B-C).

## Discussion

This work presents a bioinformatic toolset, a confirmatory study, and a web portal focused on SARS-CoV-2 genetic drift and a framework to dissect viral genomic variability by focusing on single mutations. Our analysis revealed the strong emergence of one RBD variant, possibly crucial for the virus infectivity potential.

The RBD is located from aa319-541 (one of the two crystallography analysis reports it to be at aa333-526 [24,25]. However, the major component of ACE2 contact lies in the Receptor Binding Motif (RBM), located in the aa438-506 central part. Both the S477N and the second-most frequent RBD variation N439K lie in the RBM, suggesting a selective pressure on this locus (Figure 3D).

From the polarity point of view, only the N439K switch has a change from Neutral to Polar; a recent structural topology article provided an in-depth analysis of the Energy free change of the most frequent single and clustered mutations, showing that the N439K has a strong increase of Binding Free Energy (BFE)[26]. The S477N has only a slight BFE increase that does not seem to reflect its relative increase in frequency.

Another critical mutational effect is the putative change in antibody affinity. N439K was recently reported to reduce affinity to one of the described antibodies, H00S022, while no data is still available for S477N [27].

As a limitation, we acknowledge that the consensus sequence generated by our workflow does not represent any particular clade nor viral isolate and does not take into account linkage and clustering among variations. However, the focus on specific mutational events can enable easier constant tracking for a virus that is undergoing millions of replications for clinical severity and vaccine efficacy monitoring.

## Conclusion

Tracking down and monitoring SARS-CoV-2 genomic evolution has a dramatic impact on disease severity and vaccine efficacy, even if these two variables are influenced by many factors, such as individual patient′s clinical and genetic status. Nonetheless, these novel tools can create a framework to deal with the next viral pandemic waves.

## Supporting information

manuscript tables

## List of abbreviations

*COVID-19*: *COronaVIrus Disease 2019;*
*HLA*: *Human Leukocyte Antigen;*
*MHC*: *Major Histocompatibility Complex;*
*NGS*: *Next-Generation Sequencing;*
*RBD*: *Receptor-Binding Domain;*
*RBM*: *Receptor-Binding Motif;*
*SARS-CoV-2*: *Severe Acute Respiratory Syndrome Coronavirus 2;*
*VCF*: *Variant Call Format;*
*(IEDB)*: *Immune Epitope Database and analysis resource*

## Declarations

### Ethics approval and consent to participate

Not applicable.

### Consent for publication

Not applicable.

### Availability of data and material

All R and Bash/shell scripts developed are available at https://gitlab.com/bioinfo-ire-release/covid-miner along with test files and a walkthrough.

## Acknowledgments

Not applicable.

## Authors’ contributions

Conceptualization, MP, GC, LA.; Methodology, MP, AM.; Software, AS, LDA.; Validation, LDA, ES .; Formal Analysis, MP, LDA, AM, ES.; Writing – Original Draft Preparation, MP.; Writing – Review & Editing, MP, FP, GC.; Visualization, AM, LDA,ES.; Supervision, MP;

## Figure and Table Legends

Table S1: Full list of annotated variants found in the GISAID dataset.

Table S2: Matrix of variant-country association, counts normalized by *mutations per thousand isolates*.

Table S3: Epitope prediction binding via NetMHC4 analysis.

Table S4: Epitope mutation fold-change ranking via IEDB analysis.

Table S4: BepiPred results of B-cell epitope prediction for Wild Type, S477N and N439K Spike sequences.

